# Discovery of steroidal alkaloid metabolites and their accumulation in pigs after short-term tomato consumption

**DOI:** 10.1101/2024.02.05.579005

**Authors:** Maria J. Sholola, Mallory L. Goggans, Michael P. Dzakovich, David M. Francis, Sheila K. Jacobi, Jessica L. Cooperstone

## Abstract

**Scope:** Whole tomato consumption has been shown to be more effective than lycopene alone against chronic disease risks, suggesting other phytochemicals play a role in the health properties of tomato-rich diets. Recently, metabolites of tomato steroidal alkaloids, an understudied class of secondary plant compounds, have been found in plasma, tissues, and urine. However, a comprehensive, targeted analysis to determine which steroidal alkaloid metabolites are present after tomato consumption is lacking. This study profiles and quantifies tomato steroidal alkaloids in blood for the first time.

**Methods and results:** In a two-week parallel-arm study, piglets (n = 20) were fed diets containing 10% tomato powder or a macronutrient-matched control. Steroidal alkaloids were extracted from plasma and quantified using LC-MS. Tomatidine and alpha-tomatine were detected in plasma and confirmed with standards, while mass fragmentation spectra aided in identifying 31 additional metabolites representing 9 unique masses. Concentrations averaged to 107.7 nmol/L plasma, comprising of phase I (66%) and phase II (4.5%) metabolites.

**Conclusion:** These results describe the profile and concentration of steroidal alkaloid metabolites in pig plasma after short-term tomato consumption. Our methodology and findings allow for future investigations of tomato steroidal alkaloid bioactivity using physiologically appropriate levels.

## 1. INTRODUCTION

Tomato consumption is associated with reduced risks for heart disease and various cancers, such as prostate and lung cancer ^[1–4]^. Past research has focused on lycopene when investigating the health benefits of tomatoes, as its primary source is from tomato products. It has been demonstrated to exert antioxidant potential *in vitro* ^[5]^, and epidemiological studies suggest positive health associations with higher blood levels of lycopene ^[6,7]^. However, lycopene is one of hundreds of different phytochemicals present in tomatoes, and some studies have shown that consuming whole tomato is more beneficial to health than consumption of the purified pigment ^[8,9]^. In a skin cancer mouse model, our group found that regardless of the amount of lycopene absorbed into the skin from two different tomatoes (one offering a more bioavailable form of lycopene), the same protective effect occurred, reducing skin tumors in mice by about 50% ^[10]^. This implies additional bioactive compounds contribute to the health benefits of tomatoes.

Another class of phytochemicals that accumulate in tomatoes are steroidal glycoalkaloids, which are only produced by solanaceous plants. Steroidal alkaloids are nitrogenous and cholesterol derived compounds, some of which are only found in tomatoes. The depiction of alpha-tomatine (**Error! Reference source not found**.) illustrates the common backbone (rings A-F) shared amongst all tomato steroidal alkaloids. Diversity in structures and isomers can arise from the presence of a lycotetraose (or other sugars) at position C3, F-ring modifications (e.g., acetoxylation, hydroxylation, or glycosylation on C22-27 positions), and one unsaturation of the B-ring at C5:6 ^[11,12]^.

Steroidal alkaloids are important for plant health as they confer resistance to fungal infection and herbivory ^[13,14]^, and more recent work has suggested they may also exhibit bioactive properties for humans. *In vitro* experiments have shown alpha-tomatine (the most well studied tomato steroidal alkaloid) can inhibit the growth of multiple cancer cell types including prostate, breast, colon, and liver ^[15–18]^. Studies that fed alpha-tomatine and green tomatoes (high in alpha-tomatine content) to hamsters have resulted in increased excretion of cholesterol ^[19,20]^. When tomatidine (the aglycone form of alpha-tomatine) was injected intraperitoneally in mice prior to an asthma attack-like challenge, the mice experienced lessened symptoms than those who received no treatment ^[21]^. Tomatidine has also been shown to inhibit and reduce muscle atrophy in cell lines and mice ^[22]^. More recently, it was found that tomatidine inhibited pancreatic tumor growth both *in vitro* and *in vivo* ^[23]^. These findings demonstrate that tomato steroidal alkaloids may contribute to the bioactivity of tomatoes after consumption.

Tomato steroidal glycoalkaloids and their metabolites have been recently discovered *in vivo* via metabolomics after tomato consumption in both animals ^[10,24,25]^ and humans ^[26,27]^, across plasma ^[24]^, skin ^[10]^, liver ^[25]^, and urine ^[26,27]^. These studies find different steroidal alkaloids *in vivo* as compared to what has been reported *in planta*. While glycosylated alkaloids predominate in tomato, *in vivo* metabolites are all deglycosylated products which are hydroxylated, dehydrogenated, glucuronidated or sulfonated. One study found steroidal alkaloid metabolites are excreted in urine maximally 8-16 hours after tomato consumption ^[26]^, but beyond this, very little information regarding absorption, metabolism, or excretion of steroidal alkaloids exist. It is crucial to identify metabolites present in the body, given that metabolic processes can alter compound structures, consequently influencing their bioactivity^[28]^.

The goal of this study was to comprehensively profile and quantitate steroidal alkaloid metabolites after short term consumption of tomatoes in pigs, a physiologically relevant model of human metabolism. Based on *in vivo* reports, we hypothesized tomato steroidal glycoalkaloids would undergo hydrolysis of sugars before subsequent phase I and phase II metabolism. A priori hypothesized metabolite masses in blood were explored with liquid chromatography interfaced with high resolution mass spectrometry with and without fragmentation. Compounds that met the following four criteria were considered a steroidal alkaloid metabolite if they:

1)had a ∼10 ppm accurate mass match to a proposed metabolite and at least one ^13^C isotopomer,

2)were present in plasma of tomato fed animals and absent in controls,

3)had a plausible retention for the proposed compound (with alpha-tomatine, tomatidine, dehydrotomatine, and dehydrotomatidine serving as benchmarks), and

4)had MS/MS spectra consistent with standards and previous literature reports. Annotated metabolites were quantified against authentic standards of alpha-tomatine (for glycosides) and tomatidine (for aglycones) which were spiked into pig plasma to create an in-matrix calibration curve (with alpha-solanine and solanidine as internal standards) using LC-MS.

## 2. EXPERIMENTAL SECTION

### 2.1 Experimental diet production

Roma-style processing tomatoes (*Solanum lycopersicum*) were grown to create tomato powder in this study, as described previously ^[29]^. Tomatoes derived from OH8245 × OH8243 ^[30,31]^ were grown using conventional horticultural practices, harvested using a large-scale tomato harvester and sorted for ripe fruits only. Fruits were washed and diced. Frozen tomatoes were freeze-dried and ground into a fine powder using a vertical chopper mixer, as done previously ^[10]^. Tomato powder was vacuum sealed and stored at -20 °C until use.

Diets were prepared as described in Goggans et al^[29]^. To create the control diet, a basal diet was supplemented with milk protein isolate, powdered sugar, pectin and cellulose to create a macronutrient match to the tomato powder ^[29]^. Basal diet contents can be found in **Supplementary Tables S1**. Control diet was added at 10% on a dry weight basis to basal diet. The tomato-supplemented diet was composed of basal diet supplemented with 10% tomato powder on a dry weight basis.

### 2.2 Pig study design

Male pigs (n = 20) born to six mothers were randomized to consume the control or tomato diet ^[29]^. This design is described in more detail in Goggans et al^[29]^. A graphical representation of the study design can be seen in **Figure 2**.

**Figure 1.**
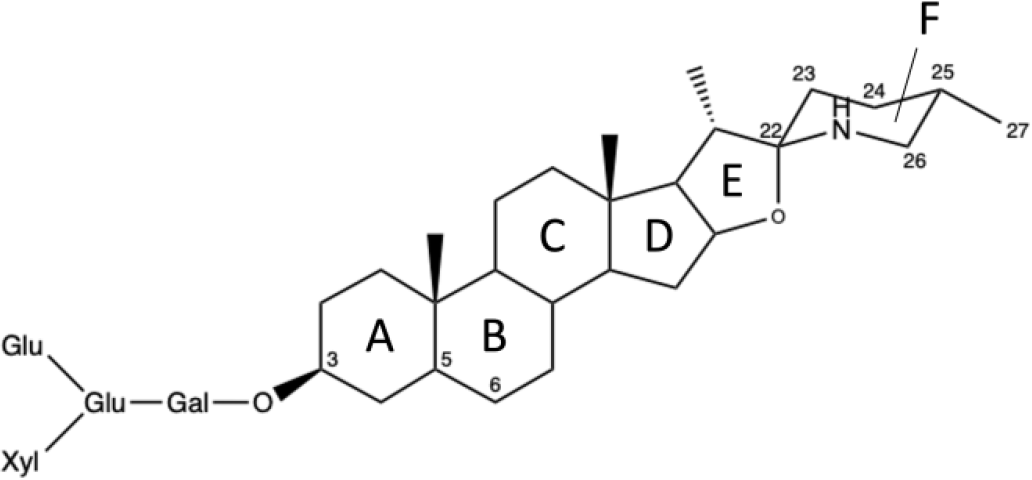
Chemical structure of alpha-tomatine with rings labeled A-F. *Glu: glucose; gal: galactose; xyl: xylose*

**Figure 2.**
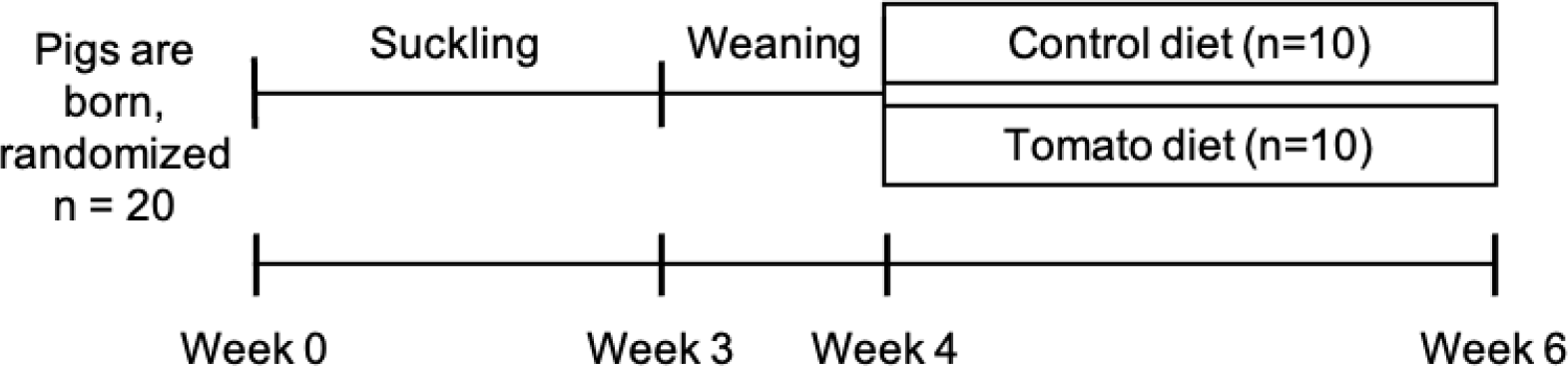
Graphical representation of study design (Goggans et al., 2022).

### 2.3 Sample Collection

After 2 weeks of dietary intervention, when pigs were 6 weeks old, they were euthanized for sample collection by captive bolt stunning. Blood was immediately drained from the body via exsanguination of the jugular vein. Blood samples were kept on ice and centrifuged at 3000 × *g* to separate blood plasma, aliquoted and frozen at -80 °C until analysis. Pools of plasma from pigs fed control and tomato diets were created separately for method development.

### 2.4 Chemicals

LC-MS Optima grade water, acetonitrile, methanol, and formic acid were all purchased from Fisher Scientific (Waltham, MA, United States). Alpha-tomatine (≥ 90%) and solanidine (≥99%) standards were purchased from Extrasynthese (Genay, France). Tomatidine HCl (≥ 95%) and alpha-solanine (≥95%) standards were purchased from Sigma-Aldrich (St. Louis, MO, United States).

### 2.5 Tomato steroidal alkaloid analysis in pig diets

#### 2.5.1 Extraction of tomato steroidal alkaloids from diet

The experimental diets were analyzed for tomato steroidal alkaloid contents. Eight technical replicates of extraction and analysis were completed to account for any potential within-sample variation. The diets were extracted as published previously ^[11]^. Five grams of pig feed were weighed into tubes containing 3/8” x 7/8” angled ceramic cutting stones (Raytech Industries, Inc. Middletown, CT, Item No. 41316). A 100 μL spike solution of two potato steroidal alkaloids, alpha-solanine and solanidine, was added as internal standards. Fifteen mL of methanol was added to the tube for extraction, along with 5 mL of water to account for typical water content of fresh tomatoes (as the extraction method was developed for tomato fruits). The samples were then homogenized in solution in a GenoGrinder 2010 (SPEX Sample Prep: Metuchen, NJ, United States) for 5 minutes at 1400 rpm. After homogenization, tubes were centrifuged for 5 minutes at 3000 × *g*. Two mL of supernatant was transferred to glass vials and 1 mL of water was added prior to analysis by UHPLC-MS/MS.

#### 2.5.2 UHPLC-MS/MS analysis of diet

Extracts were analyzed using existing methods described by Dzakovich et al^[11]^. In brief, samples were run on a Waters (Milford, MA, United States) ultra-high performance liquid chromatography (UHPLC) Acquity H-Class system interfaced with a triple quadrupole detector using multiple reaction monitoring (MRM). Analytes were separated using a C18 bridged ethylene hybrid (BEH) column (2.1 x 100 mm, 1.7 μm particle size, Waters) with a gradient of water and acetonitrile acidified with 0.1% formic acid at a flow of 0.4 mL/min. Steroidal alkaloids were quantified using external standard curves of alpha-tomatine (for glycosylated steroidal alkaloids) and tomatidine (for steroidal alkaloid aglycones), and relative to a constant quantity internal standards of alpha-solanine and solanidine, respectively. Adjustments were obtained for all compounds of interest by dividing analyte area by the corresponding internal standard area.

### 2.6 Tomato steroidal alkaloids in pig plasma

#### 2.6.1 Extraction of tomato steroidal alkaloids

The blood plasma extraction method was adapted from Cichon et al^[32]^. After thawing at room temperature, 100 μL of plasma was added to centrifuge tubes. 500 μL of cold methanol was added to each sample to precipitate proteins and extract alkaloids. Samples were homogenized in a GenoGrinder for 30 seconds at 1400 rpm and then centrifuged at 21,100 × *g* for 3 minutes. The supernatant was removed and transferred to a new microcentrifuge tube for drying. These extracts were dried down in a vacufuge (Eppendorf AG, Hamburg, Germany) at room temperature and either stored in -80 °C until analysis, or reconstituted and run immediately.

To reconstitute, 50 μL methanol was added to each sample. Samples were briefly vortexed and sonicated to ensure solubilization of the extract. Then 50 μL of water was added to each sample, and vortexed and sonicated again until dissolved. Before analysis, extracted samples were spiked with an internal standard solution of alpha-solanine and solanidine in 50% methanol and vortexed to promote integration into plasma matrix. Samples contained 0.028 pmol alpha-solanine and 0.015 pmol solanidine. For method development and creation of an in-matrix calibration curve, tomato-fed and control-fed pig plasma samples, respectively, were separately pooled and extracted following the same process. Extracts were then qualitatively and quantitatively assessed using UHPLC-QTOF-MS (for all analytes without authentic standards) and UHPLC-MS/MS (for tomatidine and alpha-tomatine).

#### 2.6.2 Metabolite discovery

Non-essential compounds undergo metabolic processes for timely excretion from the body. Using the profile of steroidal alkaloids present in the tomato diet along with a priori knowledge of mammalian metabolism of xenobiotic compounds was used to determine a list of functional group additions and other reaction products that could be plausible. The expected accurate masses resulting from different combinations of phase I and II metabolism types (**Table 1**) were calculated for each steroidal alkaloid found in tomato-supplemented feed (e.g., alpha-tomatine, hydroxytomatine, esculeoside A, lycoperosides F/G, etc.). Alpha-tomatine, for example, can be deglycosylated to form tomatidine (*m/z* 416.3528 [M+H]) which could then undergo multiple phase I hydroxylations including one (+ 18 *m/z*), two (+ 36 *m/z*), or three hydroxylations (+ 42 m/z). These hydroxyls become available sites for sulfonation and glucuronidation. Thus, for all possible hydroxylated versions of tomatidine, various combinations of phase II conjugations were considered. These included but were not limited to: 1 sulfonate addition (+ 80 *m/z*), 1 glucuronide addition (+176 *m/z*), and both sulfonate and glucuronide addition (+ 256 *m/z*). This process was repeated for the remaining steroidal alkaloids detected in tomato feed. In several instances, steroidal alkaloids from the tomato diet had overlapping theoretical metabolites (e.g., acetoxyhydroxytomatidine derived from lycoperosides F and G, esculeogenin A, acetoxytomatine, alpha-tomatine, hydroxytomatine, and tomatidine).

**Table 1.** Expected metabolic reactions for tomato steroidal alkaloids and the resulting mass difference.

| Metabolism reaction type | Reaction | Change in $m/z$ |
| --- | --- | --- |
| Phase I | Dehydrogenation | -2 |
| Phase I | Hydroxylation | +18 |
| Phase I | Acetoxylation | +60 |
| Phase II | Sulfonation | +80 |
| Phase II | Glucuronidation | +176 |

The masses calculated from these metabolic reactions were extracted from chromatograms acquired in full scan mode.

Peaks were considered putative steroidal alkaloid metabolites if they: 1) had an accurate mass within 10 ppm of the hypothesized mass and an isotope packet confirming the presence of at least one ^13^C isotopomer, 2) had a plausible retention time based on their structure relative to alpha-tomatine, dehydrotomatine, tomatidine, and dehydrotomatidine, 3) were present in the plasma of tomato fed pigs, and absent in those on control diets. Metabolites passing these criteria were then subjected to targeted MS/MS, and if they 4) had spectral characteristics similar to steroidal alkaloid standards or previously reported steroidal alkaloids in the literature^[10,24,25,33–37]^, they were level 2 ^[38]^ identified as steroidal alkaloid metabolites (**Table 4**).

##### 2.6.2.1 Full scan LC-QTOF-MS analysis

The first analyses of the blood plasma extracts were conducting using UHPLC coupled with quadrupole time-of-flight MS (UHPLC-QTOF-MS) to allow for discovery of putative metabolites. Samples were analyzed on an Agilent 1290 Infinity II UHPLC (following the same chromatographic method for diet analysis) interfaced with a 6545 QTOF-MS in positive ion mode using an electrospray ionization (ESI) source. Full scan acquisition of data was collected from a range of 100 *m/z* to 1700 *m/z* with a gas temperature of 350 °C, gas flow of 10 L/min, nebulizer gas flow at 35 psi, sheath gas flow of 11 L/min and temperature of 375 °C. An extracted ion chromatogram (EIC) was obtained for each hypothesized metabolite mass from a pool of the tomato-fed and control-fed animals. These chromatograms were then used to determine potential presence or absence of the compound.

##### 2.6.2.2 Targeted LC-QTOF-MS/MS

The same LC-QTOF-MS instrument and parameters mentioned in **Section 2.6.2.1** were used. A pre-determined list of masses was selected and fragmented with a collision energy of 30 eV. The resulting spectra was then analyzed for fragments which may have been captured during MS/MS. If at least three fragments characteristic of steroidal alkaloids were present (e.g., 161, 253, 271), the compound was assigned a putative identification. Alpha-tomatine, tomatidine, dehydrotomatine, and dehydrotomatidine peaks were analyzed in the same manner as samples for comparison. Literature MS/MS spectra was also used for identification ^[33,34]^.

### 2.6.3 Quantitative analysis of plasma extracts

Plasma steroidal alkaloids were quantified on LC-QTOF-MS or LC-MS/MS. The chromatography methods for both instruments were the same and as described in **Section 2.5.2**. For analytes that were commercially available, analysis was carried out on a triple quadrupole LC-MS/MS. The MRM parameters for each analyte (including internal standards) can be found in **Table 2**. The remaining analytes (without commercially available standards) were quantified on UHPLC-QTOF-MS with the same LC and MS parameters in **Section 2.6.2.1**. Metabolites without standards were quantified using tomatidine equivalents (as all were aglycones) and an in-matrix (i.e., plasma) calibration curve. All results produced from both LC-MS/MS and LC-MS analyses were normalized to internal standards with structural similarity (alpha-solanine and solanidine for glycoside and aglycone analytes, respectively).

**Table 2.** Parameters selected for multiple reaction monitoring MS/MS for analysis of tomato steroidal alkaloids and internal standards in plasma.

| Analyte | Retention time (min) | Precursor ion [M+H]<sup>+</sup> | Product ions [M+H]<sup>+</sup> | Cone voltage (V) | Collision energy (eV) |
| --- | --- | --- | --- | --- | --- |
| Alpha-tomatine | 5.66 | 1034.14 | 85.1*, 161 | 80 | 65, 65 |
| Tomatidine | 7.35 | 416.3 | 161.12, 255.11* | 50 | 35, 35 |
| Alpha-solanine | 5.75 | 868.27 | 98*, 398 | 90 | 85, 70 |
| Solanidine | 7.29 | 398.3 | 98.09*, 56.06 | 70 | 45, 80 |

\*Quantifying ions; other ions were used for qualitative purposes.

## 3 RESULTS AND DISCUSSION

### 3.1 Lycoperosides F/G, esculeoside A, alpha-tomatine and hydroxytomatine are the predominant steroidal alkaloids in the diet

Error! Reference source not found. lists the mean concentration of individual and total steroidal alkaloids observed in the tomato supplemented diet. There were no steroidal glycoalkaloids detected in the control diet, as no tomato products were used in that feed. Some alkaloids (e.g., lycoperosides F, G, and esculeoside A) are grouped together because they are positional or stereo-isomers, and the distinction between these isobaric compounds cannot be made here. All peak areas for isobaric masses were added together and reported as sum concentration of all the isomers.

In the diet used for this study, only 0.01% of steroidal alkaloids detected exist as aglycones; typical of steroidal alkaloids in tomatoes (both fresh and processed) which are present mostly in a glycosylated form. The most abundant steroidal alkaloid found in the tomato-supplemented diets was lycoperoside F, G, and/or esculeoside A, comprising ∼41% of total steroidal alkaloids. Hydroxytomatine and alpha-tomatine were the next most abundant, each contributing ∼25% of total alkaloid content. Together, these alkaloids combined make up 91% of the total alkaloid content in this diet. The dehydro products (i.e., dehydrotomatine, dehydrolycoperoside F/G/dehydroescueloside A) are also minor components of the diet, representing 1.7% of total alkaloids.

The diets in this study were created using raw, commercial processing tomatoes, which were freeze-dried without undergoing any heat processing. As a result, alkaloid profile observed here aligns with what has been previously reported for raw, cultivated red tomatoes ^[11,39]^. In contrast, surveys of thermally processed tomato products report an absence of lycoperoside F, lycoperoside G, or esculeoside A^[40]^, even though this is our predominant alkaloid. Thermal processing has been shown to decrease the concentration of lycoperosides F, G, and/or esculeoside A by 15-17 times ^[41]^, indicating this group of alkaloids is susceptible to heat modification. What degradation products are for this alkaloid or how they are formed are not known. Processed tomato products have been reported to consist of relatively high levels of alpha-tomatine ^[40]^ or esculeoside B ^[34,42]^ when compared to fresh tomatoes. Furthermore, esculeoside A is reported to undergo structural changes such as F-ring rearrangement, with ∼30% converting into esculeoside B after refluxing with water for 6.5 hours ^[43]^, suggesting thermal degradation of esculeoside A results in the formation of esculeoside B.

### 3.2 Tomato steroidal glycoalkaloids are metabolized into Phase I and II aglycones after consumption

Steroidal alkaloids observed in pig plasma passing all criteria mentioned in **Section 2.6.2** can be found in **Table 4** grouped by analyte. Full fragmentation data associated with every isomeric peak can be found in **Supplementary Tables S2**. In total, 9 masses and 31 peaks were annotated as tomato steroidal alkaloids or their metabolites. Ions noted as tomato steroidal alkaloid fragments are those that were abundant and matched an expected fragment characteristic of tomato steroidal alkaloids (e.g., *m/z* 161, 253, 255, 271, 273, 289, 398 ^[11,32,33,44,45]^ in positive ion mode). Fragmentation data for tomatidine and alpha-tomatine are included as these alkaloids have commercially available standards. Multiple peaks matching the selection criteria in EICs for several masses (**Supplementary Tables S2**) indicate isomeric structures. As with glycoalkaloids, some aglycone alkaloids are isobaric (e.g. lycoperoside F and esculeoside A) and cannot be distinguished without authentic standards (which are not commercially available). When multiple peaks were discovered in a chromatogram for one mass, each peak was scrutinized separately following the same selection criteria. In many instances, several corresponding peaks of one mass had the same expected fragments of steroidal alkaloid aglycones in their mass spectra. While fragments were the same, their abundances varied, inferring that there might be multiple positional isomers for some masses.

Tomato steroidal alkaloids have been found to exist as aglycones in a variety of tissues in mice ^[10,25,32]^, and in human urine ^[26]^. Based on the glycoalkaloids observed in tomato-supplemented diets, we hypothesized that the metabolites we would find in plasma would have the backbone of the following aglycones: tomatidine, acetoxytomatidine, dehydrotomatidine, and esculeogenin B, before phase I/II processes. This result is demonstrated here (apart from alpha-tomatine), supporting evidence that tomato steroidal glycoalkaloids are deglycosylated *in vivo* before absorption. We hypothesize the cleavage of sugars occurs in the small intestines by beta-glucosidases expressed in epithelial cells, as seen with other plant glycosides like flavonoids^[46]^. Interestingly, it was found in an *ex vivo* pig cecum model that alpha-tomatine underwent minimal degradation after incubation with cecum suspension^[47]^, suggesting that gut microbial metabolism may also play a lesser role in the hydrolysis of alpha-tomatine and potentially other tomato steroidal alkaloids.

Desaturated alkaloids (e.g., dehydrotomatidine) were not observed in pig plasma. Considering the low concentrations of dehydrotomatine, dehydrolycoperoside F, G and dehydroesculeoside A in the experimental diet (Error! Reference source not found.), it would be expected that lower levels of metabolites derived from these compounds would be present as compared to saturated alkaloids. Given abundance of some steroidal alkaloids observed near the bottom of linear range, these compounds might exist, but under the limits of detection here. As desaturated alkaloids can result from tomato or phase I human metabolism, the lack of their observation suggests desaturation is not extensively occurring as a phase I reaction. This finding is inconsistent with previous reports of detected desaturated alkaloids in mouse plasma, skin, and liver ^[10,25,32]^. However, these studies did not analyze the steroidal glycoalkaloid content comprehensively in the diets which makes comparing dietary exposure to metabolites observed *in vivo* challenging. These aforementioned studies were carried out in mice which have different gastrointestinal physiology compared to pigs, namely in the small and large intestines. Pigs possess digestive system traits more similar to humans than mice, such as comparable ratios of small intestine length to bodyweight and similar types of epithelial cells comprising the intestines ^[48,49]^. It is possible that the absence of dehydro metabolites observed in our study could be due to dietary or organismal differences, or both.

### 3.3 Phase I metabolites of tomato steroidal alkaloids accumulate the highest in blood plasma after consumption

Here we found that tomato steroidal alkaloid metabolites accumulate to 107.7 (48.9 – 206) nmol/L plasma (Error! Reference source not found.), with phase I metabolites accounting for 66% total metabolites detected. Phase II metabolites were only sulfonate conjugates which represent 4.5% of steroidal alkaloids detected in plasma. Tomatidine accounts for 7.4% of steroidal alkaloids observed in plasma. The presence of tomatidine in plasma could be because alpha-tomatine is deglycosylated prior to (or after) absorption, and/or the small amount consumed from the tomato diet is absorbed directly. Our study does not allow the distinguishing of these two situations.

While aglycones made up majority of plasma steroidal alkaloids, alpha-tomatine was the only glycosylated alkaloid detected and accounted for 22% of alkaloids detected in plasma. The absorption of alpha-tomatine was unexpected as larger polar molecules (e.g., flavonoid glycosides) are typically poorly absorbed^[50–52]^, and its presence has not been reported in tomato-fed mice ^[10,25,32]^. Young mammals tend to have “leaky guts,” meaning the junctions between cells in their intestinal walls are not as compacted as those in adults. A leaky gut could enable passive absorption of compounds that typically do not get absorbed in adults ^[53]^. If steroidal alkaloids are enzymatically hydrolyzed at the intestinal brush border, our use of young piglets might explain the relatively large amount of alpha-tomatine entering blood circulation. This hypothesis requires additional experimental testing.

### 3.4 Positioning of hydroxyl groups on rings A-D of steroidal alkaloid backbone results in fragmentation patterns that allow for distinction between isobars

MS/MS experiments on plasma steroidal alkaloids revealed fragments *m/z* 271 and 253 for multiple precursor ions which have been reported for desaturated steroidal alkaloids ^[36,37,40,54]^. However, in our case these fragments often appeared alongside *m/z* 289. These ions (*m/z* 253, 271, and 289) each differ by one water loss – a common neutral loss observed during tandem MS for hydroxylated aliphatic compounds ^[55,56]^. Upon investigation, it was found that these fragments may correspond to breakage of the steroidal alkaloid backbone with additional hydroxyl groups located between rings A-D. This observation was used to distinguish isobaric masses putatively identified as dihydroxytomatidine and esculeogenin B (**Figure 3**).

**Figure 3.**
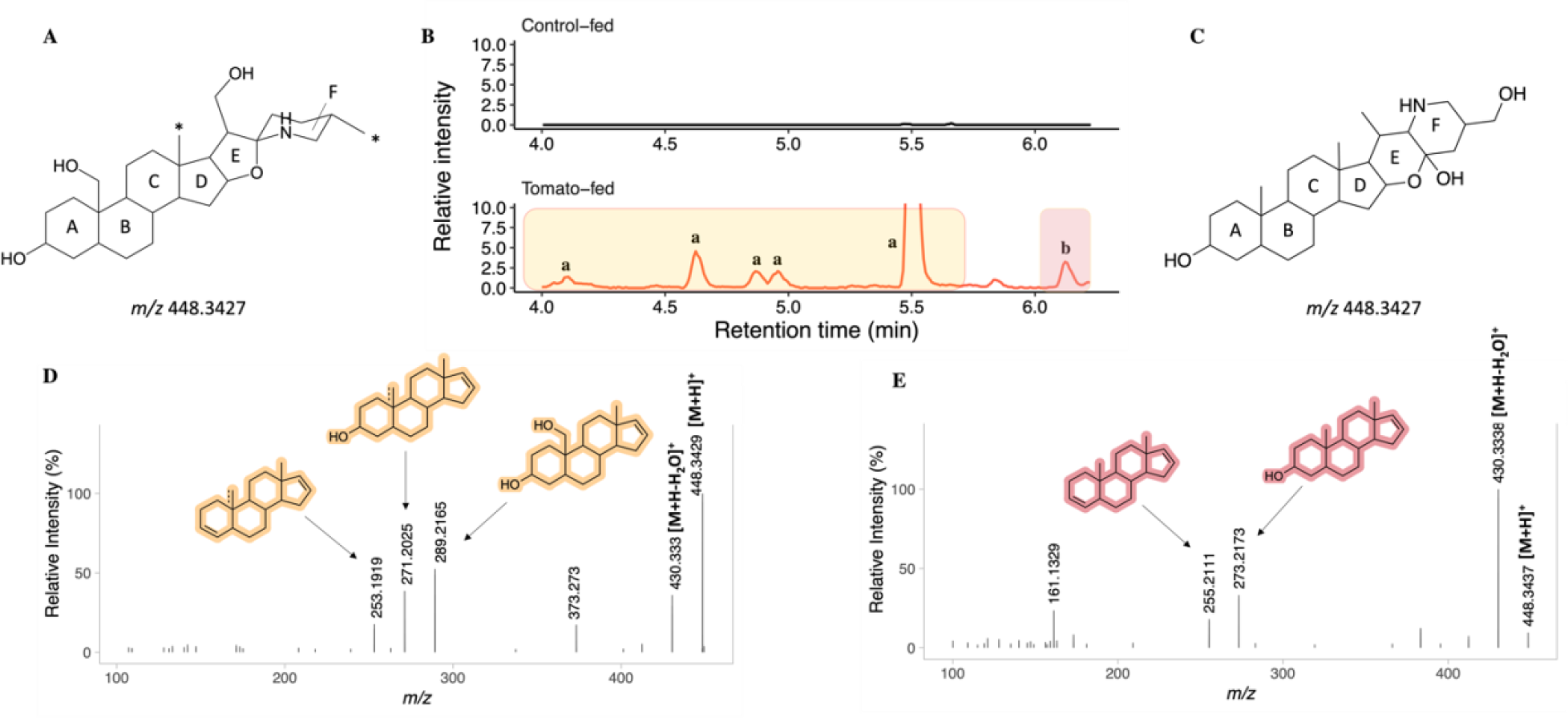
**A**) Tentative structure of dihydroxytomatidine *\*other possible hydroxyl locations*; **B**) extracted ion chromatogram of *m/z* 448 from control- and tomato-fed pig plasma samples; **C**) chemical structure of esculeogenin B; **D**) MS/MS spectrum corresponding to peaks **a**; **E**) MS/MS spectrum corresponding to peak **b**. Peak **a**: putatively identified as dihydroxytomatidine isomers; Peak **b**: a dihydroxytomatidine isomer or esculeogenin B

**Figure 3B** shows the chromatogram extracted from m/z 448, representing both dihydroxytomatidine and esculeogenin B. Several peaks appear for this mass in tomato-fed pig plasma, but not in control-fed pig plasma, indicating the presence of isomeric and/or isobaric structures from tomato ingestion. Peaks annotated **a** and **b** in **Figure 3B** exhibit MS/MS spectra characteristic of steroidal alkaloids. The structure of dihydroxytomatidine (**Figure 3A**) is tentative, due to unknown hydroxyl locations. For minimized steric hindrance, we hypothesize that configurations with one added hydroxyl between rings A-D and the other between rings E-F are more likely. Calculations based on typical steroidal alkaloid ring fragmentation^[44,45]^ and an additional hydroxyl group on rings A-D resulted in m/z 289, followed by a neutral water loss to m/z 271, followed by another water loss to form m/z 253 are illustrated in **Figure 3D**. This was the fragmentation pattern was observed for peaks **a** (**Figure 3B**). In the case that dihydroxytomatidine exists as the configuration proposed in **Figure 3A**, this compound would be representative of peaks **a**, accounting for 11.976 nmol/L steroidal alkaloids detected in plasma. See **Supplementary Figures S1** for graphical representations of proposed mass fragmentation pathways for dihydroxytomatidine isomers.

After isolating esculeoside B from tomato juice and performing acid hydrolysis, Nohara et al. synthesized esculeogenin B (**Figure 3C**) and confirmed its structure with nuclear magnetic resonance (NMR)^[42]^. Given that the confirmed structure of esculeogenin B does not contain additional hydroxyls between rings A-D, the fragmentation pattern shown in **Figure 3D** is less plausible for this compound. Peak **b** in **Figure 3B** has the MS/MS spectrum shown in **Figure 3E**, which exhibits common fragments seen for steroidal alkaloids without additional hydroxyls on rings A-D^[10,25,26,32,44]^. Peak **b** could represent esculeogenin B, although there is a possibility that dihydroxytomatidine could exist in a conformation with both hydroxyl groups on rings E-F. When cross-referenced with semi-purified standard following the same protocol, Hövelmann et al. did not detect esculeogenin B in human urine following tomato juice consumption^[26]^.

Without authentic standards, we were unable to conclusively distinguish peak **b** as dihydroxytomatidine or esculeogenin B using this rationale. In the end, whether there is any biological significance of the presence of dihydroxytomatidine vs. esculeogenin B is not known.

Two peaks representing sulfonated dihydroxytomatidine/esculeogenin B (*m/z* 528) were observed. For both peaks, fragments *m/z* 353, 273, and 255 (Error! Reference source not found., **Supplementary Tables S2**), corresponds to a sulfate loss (Δ80 Da) from 353 and a subsequent water loss (Δ18 Da) from 273. A neutral loss of *m/z* 80 is typical for sulfonate conjugates and indicates sulfate removal (-SO_3_)^[55,57,58]^. Consistent with our findings, Hövelmann et al. ^[26]^ reported tandem MS fragments *m/z* 353 and 255 for the same precursor *m/z* 528 detected in urine following tomato juice consumption. They tentatively labeled *m/z* 353 as a sulfate group attached to the A ring of the steroidal fragment *m/z* 273 since it contains the only hydroxy group available for sulfonation. This characteristic could apply to both sulfonated dihydroxytomatidine (if hydroxylation occurred on rings E-F) and sulfonated esculeogenin B.

### 3.5 The primary steroidal alkaloid metabolite mass detected in blood is tentatively identified as trihydroxytomatidine

Trihydroxytomatidine (**Figure 4A**) and/or hydroxyesculeogenin B **(Figure 4B**) isomers accounted for ∼42% of steroidal alkaloid metabolites found in blood. Several peaks present in the EIC for *m/z* 464 were present in tomato-fed plasma samples but absent in control-fed pig plasma (**Figure 4C**). MS/MS spectra corresponding to peaks in **Figure 4D** were representative of steroidal alkaloids. However, the same rationale used to distinguish dihydroxytomatidine and esculeogenin B could not be used for any peaks in this case. All peaks had MS/MS spectra with fragments hypothesized to be representative of breakage of the steroidal alkaloid ring structure with an additional hydroxyl present (m/z 289, 271, 253). Both trihydroxytomatidine and hydroxyesculeogenin B could contain a hydroxyl between rings A-D (**Figure 4A-B)**, but the lack of confirmed chemical identities for either makes drawing conclusions from these fragments challenging. Hövelmann et al. report *m/z* 464 in urine as hydroxyesculeogenin B isomers with notable MS/MS fragments 289 and 271^[26]^, lacking the additional water loss (i.e., *m/z* 253) seen in this work. Considering subjects in Hövelmann et al. consumed thermally processed tomatoes ^[26]^ (known to contain higher levels of esculeoside B compared to tomatoes), their annotation could be plausible in that case.

**Figure 4.**
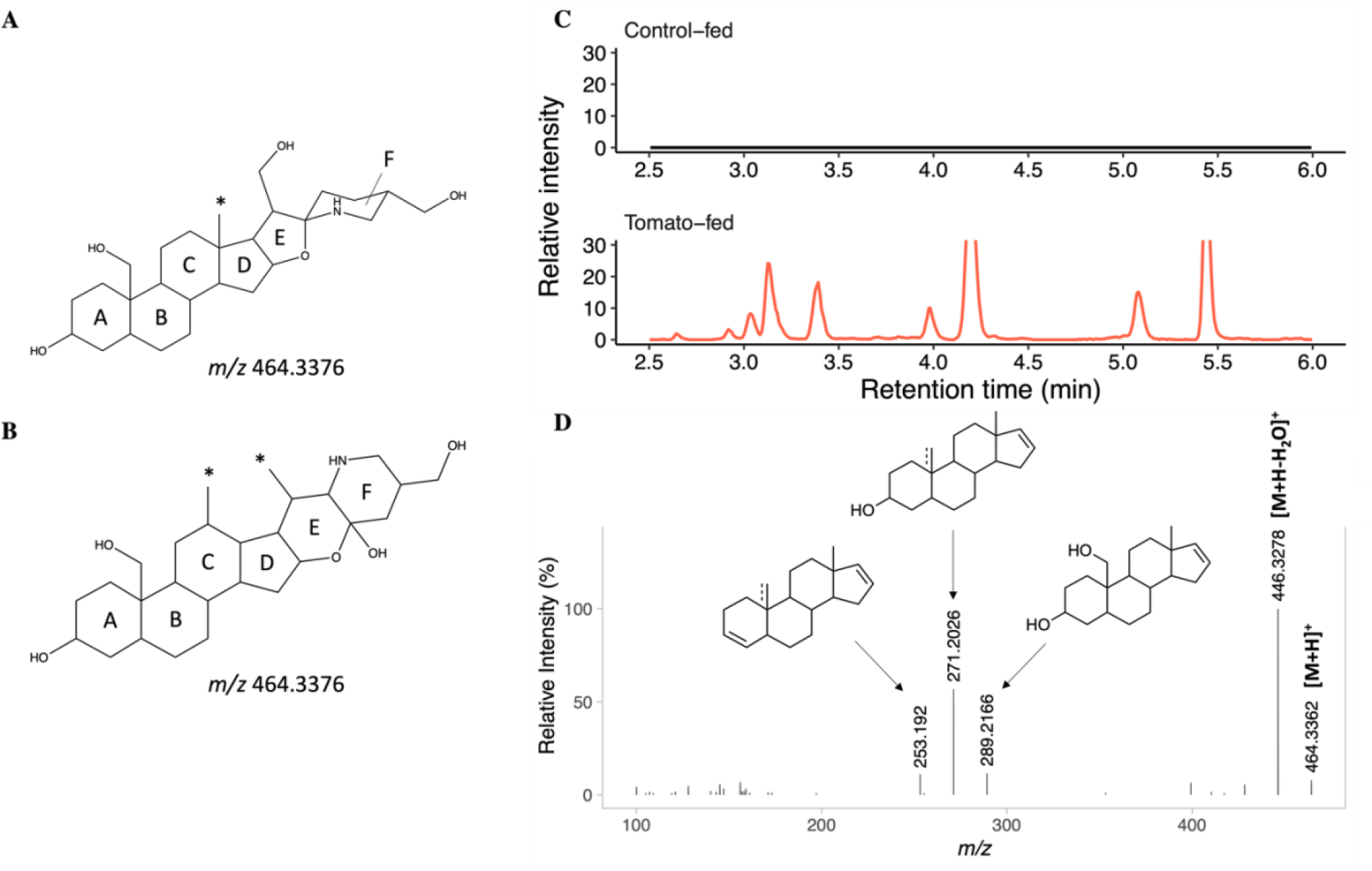
Tentative structures of A) trihydroxytomatidine and B) hydroxyesculeogenin B. *other possible hydroxyl locations; C) extracted ion chromatogram of m/z 464 from control- and tomato-fed pig plasma samples, and D) MS/MS spectrum corresponding to the 9 peaks.

**Figure 5.**
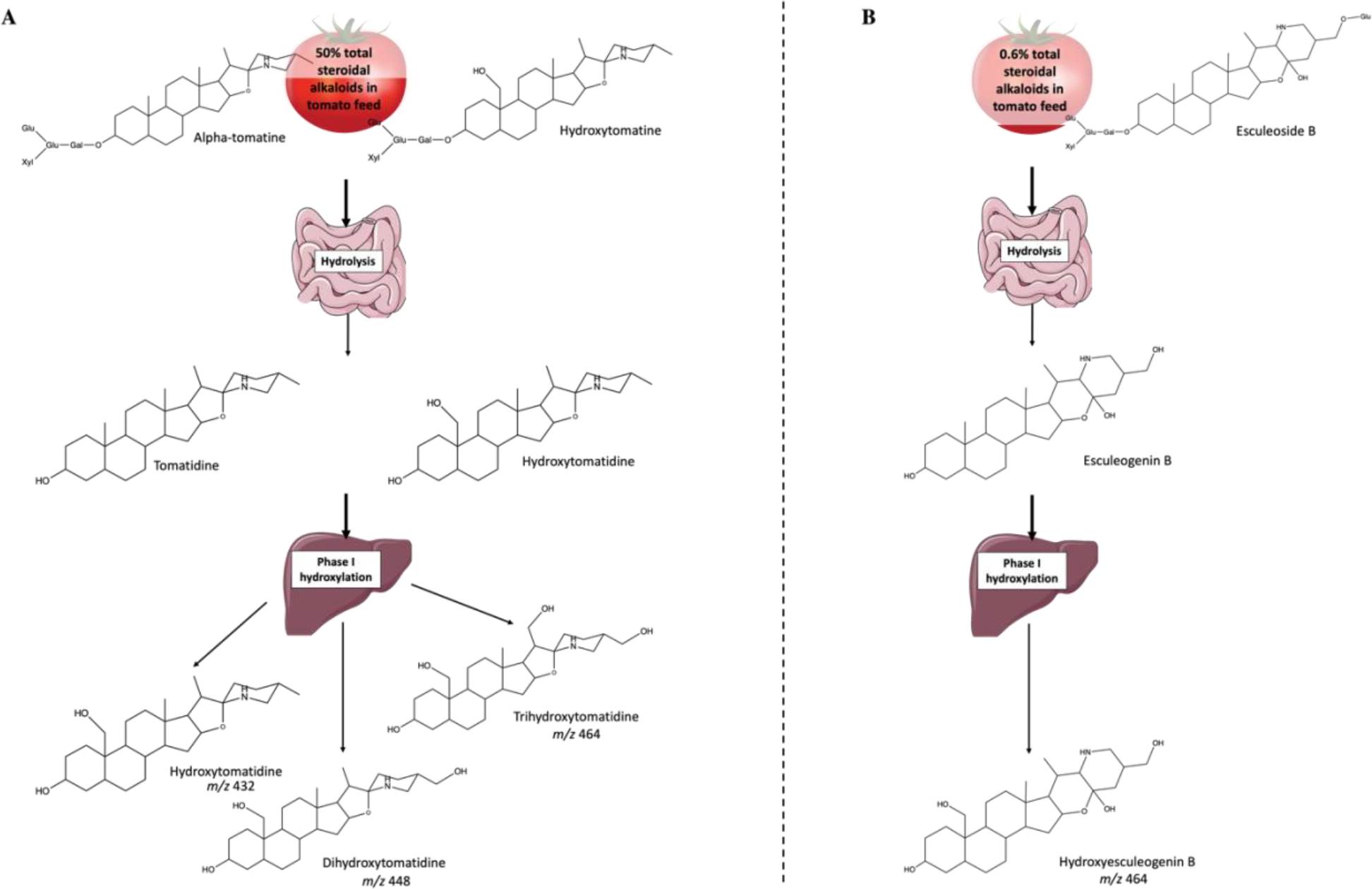
Possible metabolism pathways of A) alpha-tomatine, hydroxytomatine, and B) esculeoside B. Location of hydroxyls on hydroxytomatine, dihydroxytomatidine, trihydroxytomatidine, and hydroxyesculeogenin B are tentative.

Nonetheless, steroidal glycoalkaloid contents measured in the tomato diet for this work provides a basis for distinguishing hydroxyesculeogenin B and trihydroxytomatidine. Due to the F-ring arrangement of esculeoside B, it is the only likely precursor for hydroxyesculeogenin B. Esculeoside B is present at relatively low concentration (22.29 μg/kg) in the minimally processed tomato diet fed to pigs in our study (**Table 3**. Average and standard deviation (n = 8) of tomato steroidal glycoalkaloid levels in tomato-supplemented pig feed). Lycoperosides F/G, esculeoside A, hydroxytomatine, alpha-tomatine, acetoxytomatine, and tomatidine all represent potential precursors for the formation of trihydroxytomatidine. Collectively, these compounds account for approximately 99% of steroidal alkaloids detected in tomato feed. Acetoxylated steroidal alkaloids detected (lycoperosides F/G, esculeoside A, and acetoxytomatine) might undergo hydrolysis of acetoxy groups before subsequent hydroxylations to produce trihydroxytomatidine. However, it is also possible that acetoxy groups remain unaltered, already being oxidized enough for efficient absorption or excretion. Both cases could be true simultaneously but can’t be verified here. For the sake of clarity, we focus on alpha-tomatine and hydroxytomatine, constituting 50% of steroidal glycoalkaloids in the tomato diet, versus esculeoside B, which represents a mere 0.6% of the glycoalkaloids. Probability-wise, trihydroxytomatidine may represent more peaks corresponding to *m/*z 464 than hydroxyesculeogenin B. A simplified illustration of possible metabolism pathways can be seen in Error! Reference source not found.

**Table 3.** Average and standard deviation (n = 8) of tomato steroidal glycoalkaloid levels in tomato-supplemented pig feed.

| Analyte | Concentration (µg/kg feed) |
| --- | --- |
| Tomatidine | 0.25 ± 0.1 |
| Dehydrolycoperoside F, G, Dehydroesculeoside A | 16.58 ± 3.9 |
| Esculeoside B | 22.29 ± 2.8 |
| Dehydrotomatine | 44.05 ± 7.4 |
| Acetoxytomatine | 208.1 ± 30.8 |
| Alpha-tomatine | 875.5 ± 142 |
| Hydroxytomatine | 887.7 ± 119 |
| Lycoperoside F, G, Esculeoside A | 1405 ± 164 |
| Total steroidal alkaloids | 3459 ± 431 |

**Table 4.**
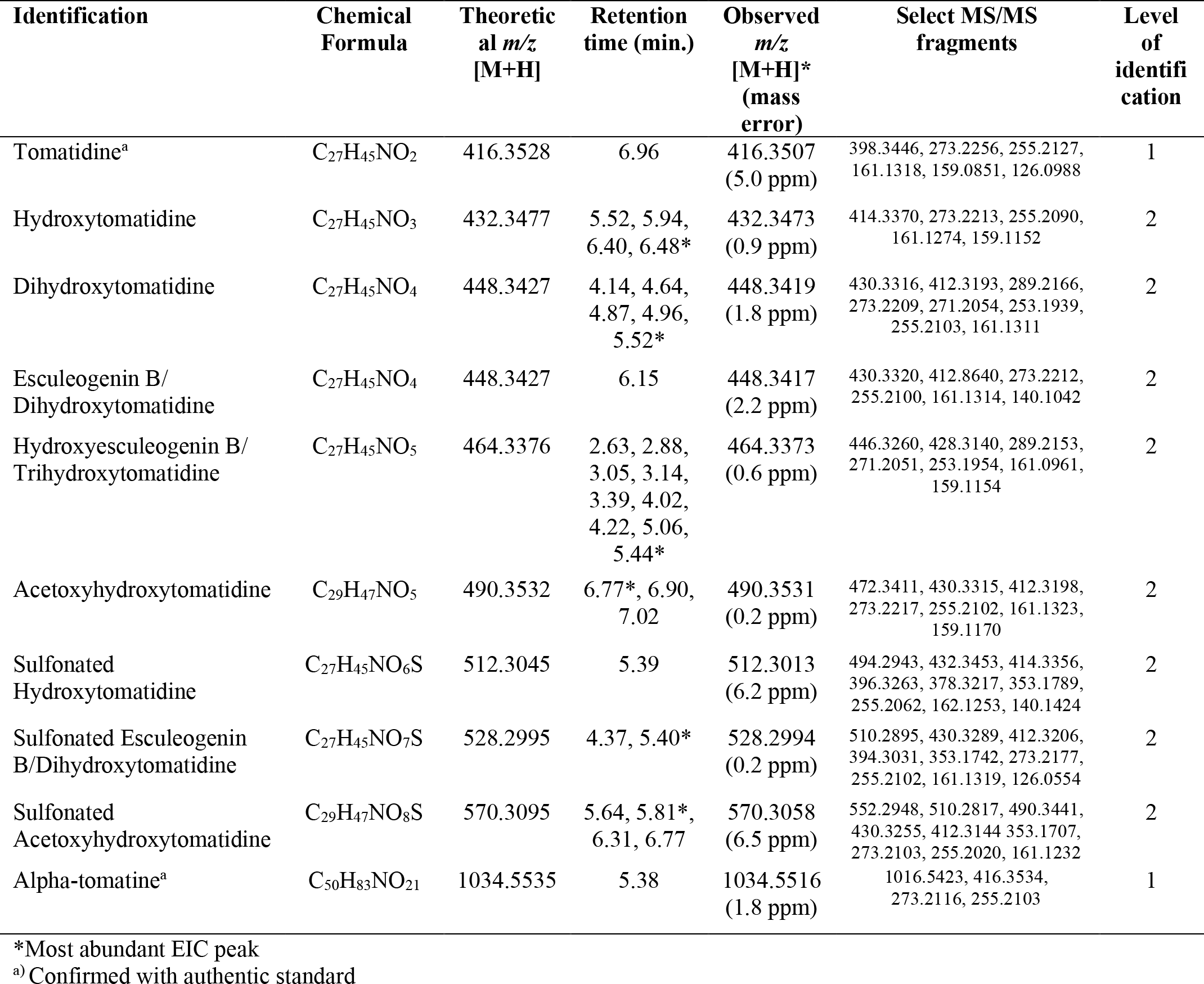
Tomato steroidal alkaloids detected in pig plasma and their corresponding fragments.

| Identification | Chemical Formula | Theoretical $m/z$ [M+H] | Retention time (min.) | Observed $m/z$ [M+H]* (mass error) | Select MS/MS fragments | Level of identification |
| --- | --- | --- | --- | --- | --- | --- |
| Tomatidine<sup>a</sup> | C<sub>27</sub>H<sub>45</sub>NO<sub>2</sub> | 416.3528 | 6.96 | 416.3507 (5.0 ppm) | 398.3446, 273.2256, 255.2127, 161.1318, 159.0851, 126.0988 | 1 |
| Hydroxytomatidine | C<sub>27</sub>H<sub>45</sub>NO<sub>3</sub> | 432.3477 | 5.52, 5.94, 6.40, 6.48* | 432.3473 (0.9 ppm) | 414.3370, 273.2213, 255.2090, 161.1274, 159.1152 | 2 |
| Dihydroxytomatidine | C<sub>27</sub>H<sub>45</sub>NO<sub>4</sub> | 448.3427 | 4.14, 4.64, 4.87, 4.96, 5.52* | 448.3419 (1.8 ppm) | 430.3316, 412.3193, 289.2166, 273.2209, 271.2054, 253.1939, 255.2103, 161.1311 | 2 |
| Esculeogenin B/<br>Dihydroxytomatidine | C<sub>27</sub>H<sub>45</sub>NO<sub>4</sub> | 448.3427 | 6.15 | 448.3417 (2.2 ppm) | 430.3320, 412.8640, 273.2212, 255.2100, 161.1314, 140.1042 | 2 |
| Hydroxyesculeogenin B/<br>Trihydroxytomatidine | C<sub>27</sub>H<sub>45</sub>NO<sub>5</sub> | 464.3376 | 2.63, 2.88, 3.05, 3.14, 3.39, 4.02, 4.22, 5.06, 5.44* | 464.3373 (0.6 ppm) | 446.3260, 428.3140, 289.2153, 271.2051, 253.1954, 161.0961, 159.1154 | 2 |
| Acetoxyhydroxytomatidine | C<sub>29</sub>H<sub>47</sub>NO<sub>5</sub> | 490.3532 | 6.77*, 6.90, 7.02 | 490.3531 (0.2 ppm) | 472.3411, 430.3315, 412.3198, 273.2217, 255.2102, 161.1323, 159.1170 | 2 |
| Sulfonated<br>Hydroxytomatidine | C<sub>27</sub>H<sub>45</sub>NO<sub>6</sub>S | 512.3045 | 5.39 | 512.3013 (6.2 ppm) | 494.2943, 432.3453, 414.3356, 396.3263, 378.3217, 353.1789, 255.2062, 162.1253, 140.1424 | 2 |
| Sulfonated Esculeogenin B/<br>Dihydroxytomatidine | C<sub>27</sub>H<sub>45</sub>NO<sub>7</sub>S | 528.2995 | 4.37, 5.40* | 528.2994 (0.2 ppm) | 510.2895, 430.3289, 412.3206, 394.3031, 353.1742, 273.2177, 255.2102, 161.1319, 126.0554 | 2 |
| Sulfonated<br>Acetoxyhydroxytomatidine | C<sub>29</sub>H<sub>47</sub>NO<sub>8</sub>S | 570.3095 | 5.64, 5.81*, 6.31, 6.77 | 570.3058 (6.5 ppm) | 552.2948, 510.2817, 490.3441, 430.3255, 412.3144, 353.1707, 273.2103, 255.2020, 161.1232 | 2 |
| Alpha-tomatine<sup>a</sup> | C<sub>50</sub>H<sub>83</sub>NO<sub>21</sub> | 1034.5535 | 5.38 | 1034.5516 (1.8 ppm) | 1016.5423, 416.3534, 273.2116, 255.2103 | 1 |

\*Most abundant EIC peak
<sup>a</sup>) Confirmed with authentic standard

**Table 5.**
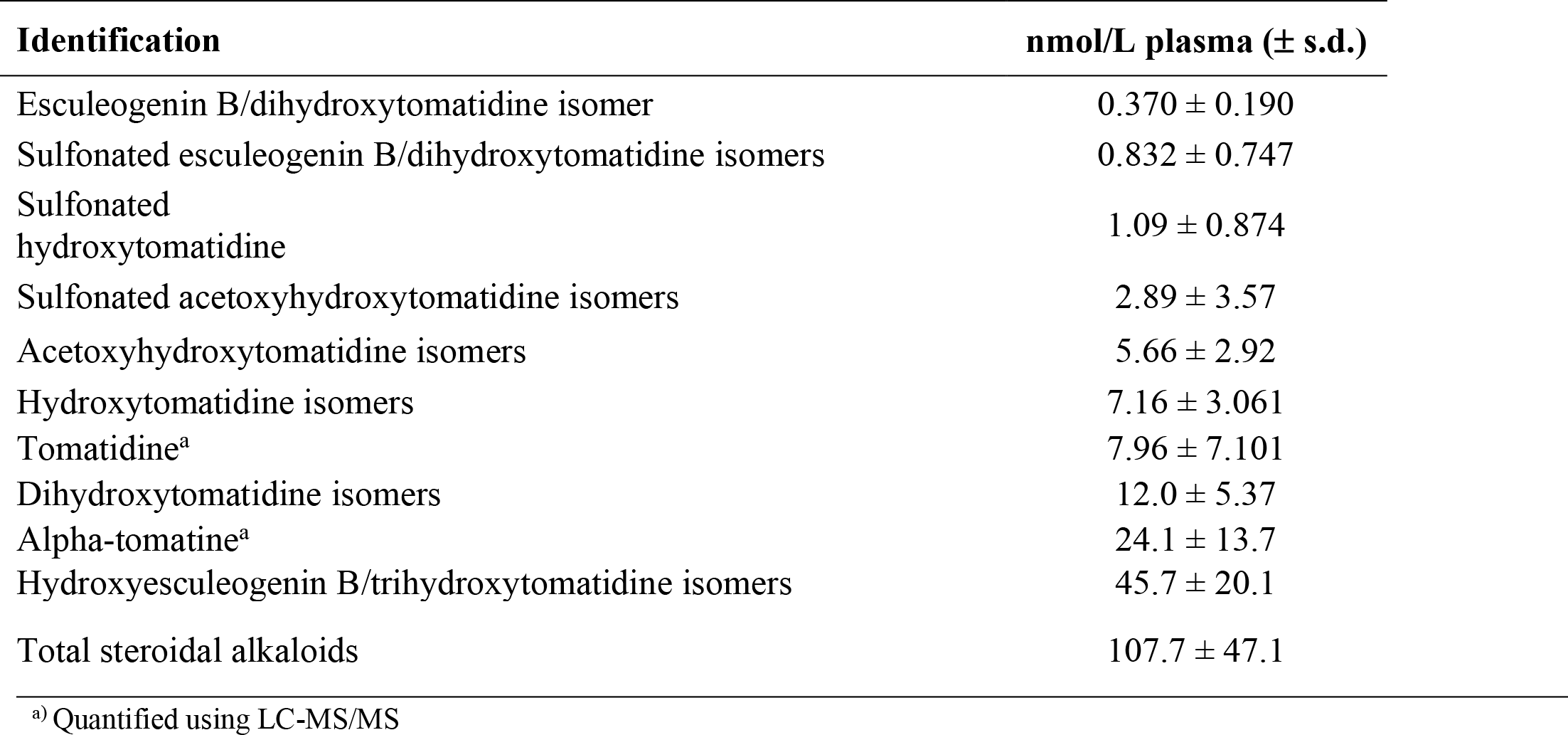
Mean plasma tomato steroidal alkaloid concentrations in pigs (n = 10) after two weeks of daily tomato consumption.

## 4 Concluding remarks

This study reports that steroidal alkaloids reach an average of 107.7 nmol/L in plasma after short term consumption by pigs, primarily as phase I and II metabolites. Phase I metabolites were the most predominant forms found in blood. Our findings align with previous reports of hydrophilic tomato steroidal alkaloid metabolites after tomato consumption. Further research needs to be pursued to fully understand the metabolic processes of tomato steroidal alkaloids. In many cases, metabolites could be derived from multiple steroidal alkaloids detected in the diet. It could not be determined here whether steroidal alkaloid aglycones were absorbed directly from the tomato diet after sugar removal, changed through phase I enzymes, or both.

The structural similarities and isomeric variations of several tomato steroidal alkaloids and their metabolites makes it a challenge to conclusively identify them. However, combining diet analysis with tandem MS provides stronger justifications for the annotation of some masses (e.g., distinguishing dihydroxytomatidine from esculeogenin B). We have rationalized a group of 3 fragments occurring from steroidal alkaloid ring breakage, which help elucidate hydroxyl configurations of isomers. To the best of our knowledge, this cluster of fragments (289, 271, 253) have only recently been reported from our group^[25]^. The discovery of new steroidal alkaloid fragmentation patterns improves accuracy of detection in both mammalian samples and plant material.

Stemming from occurrences of acute toxicity in humans after the ingestion of high levels of steroidal glycoalkaloids in greened potatoes^[59–63]^ (e.g., alpha-solanine and alpha-chaconine), there is a misconception that all steroidal alkaloids are anti-nutritional^[64,65]^. Yet, as discussed earlier, growing evidence suggests steroidal alkaloids derived from tomatoes could be health beneficial^[10,15,19,20]^. In this study, pigs were freely fed diet powders made with 10% processing-type tomatoes, like ones used for commercial consumer products. The tomato diets had an average steroidal alkaloid content of ∼3.5 mg/kg diet. For comparison, one serving size (126 g) of fresh market tomatoes from a grocery store survey consisted of an average of ∼3.4 mg steroidal alkaloids^[11]^. Thus, in our work, it is established that steroidal alkaloids are absorbed from tomatoes at normal doses.

To our knowledge, this is the first quantitative report of tomato steroidal alkaloids *in vivo*.Our accumulation data will help establish the physiological significance of these compounds and their metabolites and further assist in elucidating the health benefits of tomatoes.

