## Supplementary Figures for "Discovery of steroidal alkaloid metabolites and their accumulation in pigs after short-term tomato consumption"

**Supplementary Figure S1.** Proposed fragmentation scheme of dihydroxytomatidine after MS/MS fragmentation at 30 eV in positive electrospray ionization mode.
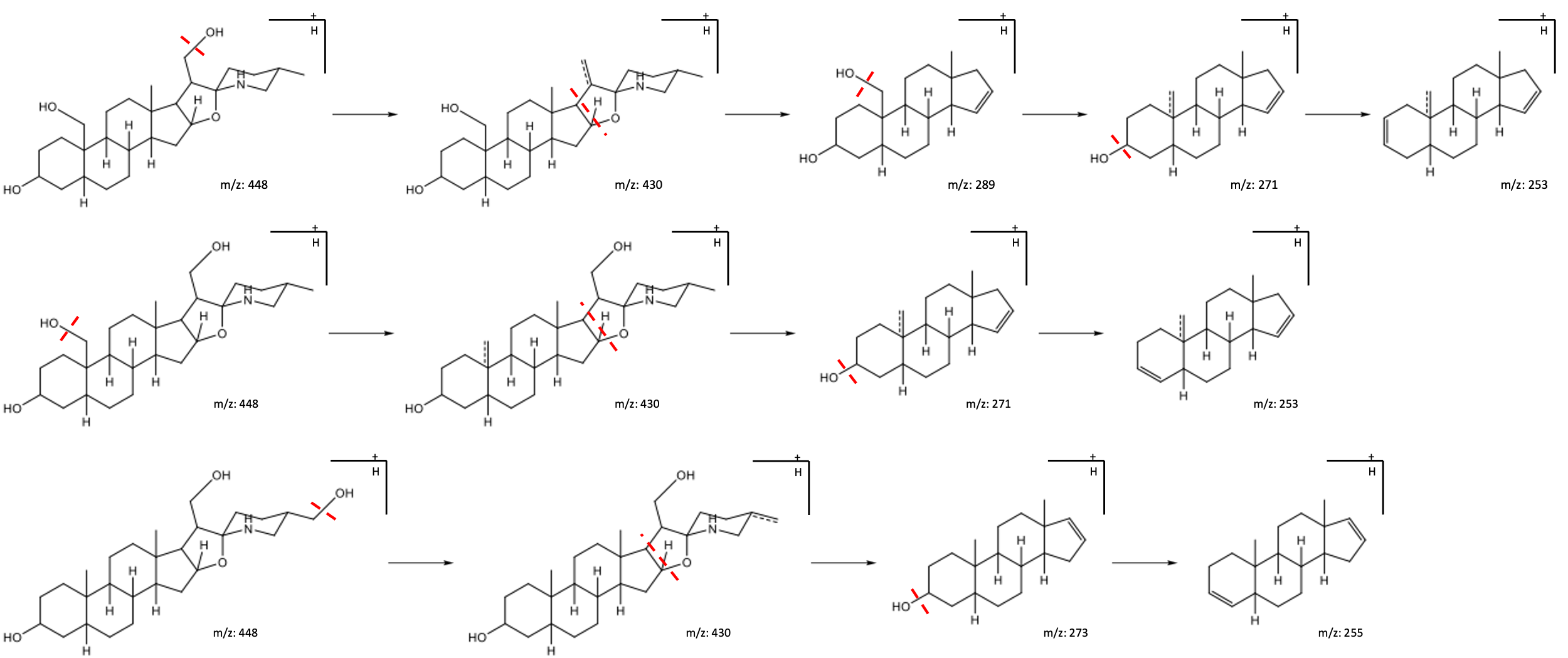
